# Testis Specific Serine Kinase 6 (TSSK6) is abnormally expressed in colorectal cancer and promotes oncogenic behaviors

**DOI:** 10.1101/2024.01.08.574658

**Authors:** Magdalena Delgado, Zachary Gallegos, Steve Stippec, Kathleen McGlynn, Melanie H. Cobb, Angelique W. Whitehurst

## Abstract

Cancer testis antigens (CTAs) are a collection of proteins whose expression is normally restricted to the gamete, but abnormally activated in a wide variety of tumors. The CTA, Testis specific serine kinase 6 (TSSK6), is essential for male fertility in mice. Functional relevance of TSSK6 to cancer, if any, has not previously been investigated. Here we find that TSSK6 is frequently anomalously expressed in colorectal cancer and patients with elevated TSSK6 expression have reduced relapse free survival. Depletion of TSSK6 from colorectal cancer cells attenuates anchorage independent growth, invasion and growth in vivo. Conversely, overexpression of TSSK6 enhances anchorage independence and invasion in vitro as well as in vivo tumor growth. Notably, ectopic expression of TSSK6 in semi-transformed human colonic epithelial cells is sufficient to confer anchorage independence and enhance invasion. In somatic cells, TSSK6 co-localizes with and enhances the formation of paxillin and tensin positive foci at the cell periphery, suggesting a function in focal adhesion formation. Importantly, TSSK6 kinase activity is essential to induce these tumorigenic behaviors. Our findings establish that TSSK6 exhibits oncogenic activity when abnormally expressed in colorectal cancer cells. Thus, TSSK6 is a previously unrecognized intervention target for therapy, which could exhibit an exceptionally broad therapeutic window.

## Introduction

Cancer testis antigens (CTAs) are a set of > 200 proteins whose expression is normally confined to the testis, oocyte and/or placenta, but frequently anomalously activated in cancer(1, 2). As the testis is an immune-privileged site, expression of these proteins in somatic cells can generate immunogenic antigens recognized by T-cells (1). Indeed T-cell mediated therapies targeting CTAs have exhibited clinical efficacy (3, 4). In addition to immunogenic properties, recent studies suggest CTAs can be functionally integrated into the tumor cell regulatory environment. Specifically, individual CTAs have been implicated in degrading tumor suppressors, enhancing oxidative phosphorylation, regulating transcriptional networks, influencing genomic integrity, and altering immune recognition (5–14). Thus, an emerging hypothesis is that CTAs are not innocent bystanders during tumorigenesis, but are intimately engaged in promoting neoplastic behaviors.

The CTA family includes Testis Specific Serine Kinase 6 (TSSK6) (15). TSSK6 is a member of a 5-protein kinase family found on the CAMK branch of the kinome (16). Deletion of TSSK6 in mice leads to male, but not female, infertility (17). As expected given TSSK6’s testes-restricted expression pattern, these mice exhibit no defects in development or adult tissues and female mice develop and reproduce normally (17). In sperm, TSSK6 is expressed post-meiotically and localizes to the nucleus in spermatids and the posterior head in spermatocytes. TSSK6-null sperm exhibit defective histone-to-protamine transition (required for post-meiotic chromatin remodeling), as well as a decrease in actin polymerization in spermatocyte (18, 19). The results are striking morphological defects in sperm including nuclear deformities, a hairpin shape with heads facing backwards, or completely detached heads (19).

TSSK6 was initially classified as a cancer-testis antigen based on its restricted mRNA expression to the testes and detection in lung cancer, sarcoma and lymphoma (15). A recent study has indicated that TSSK6 mRNA expression correlates with T-cell diversity (20). However, TSSK6 protein expression in cancer and any functional relevance to tumorigenesis have not been investigated. If TSSK6 is active when aberrantly expressed in somatic cells it could theoretically induce a tumor-specific signaling pathway that may have dramatic consequences on cellular behaviors.

Here we demonstrate that TSSK6 is frequently abnormally expressed in colorectal cancer, where it correlates with reduced relapse free survival. We find that TSSK6 supports tumor cell viability, anchorage independence, invasion and growth in vivo. Importantly, TSSK6 exhibits kinase activity when ectopically expressed and this activity is crucial for supporting these oncogenic behaviors. Mechanistically, we find that TSSK6 co-localizes with cytoskeletal regulatory proteins at focal adhesions and enhances paxillin and tensin-positive foci formation. Our findings establish TSSK6 as a novel oncogenic protein, which when abnormally expressed in somatic cells can alter signaling networks to promote anchorage independent growth, invasion and tumor growth in vivo.

## Results

### TSSK6 is frequently abnormally expressed in colorectal cancer

TSSK6 mRNA expression is normally restricted to the testes with limited expression in adult normal human tissues, including ovary (Supplemental Figure 1A). To determine if TSSK6 expression is present in human tumors, we queried the PanCancer Atlas for TSSK6 mRNA expression. Here we found that TSSK6 is elevated in ∼6% of cancers (z-score > 2.0) and expressed in a wide range of tumors types (Supplemental Figure 1B) (21). We then asked whether TSSK6 mRNA expression correlated with patient outcomes in breast, lung, ovarian, epigastric, pancreatic and colon cancer (Supplemental Figure 1C). Here, we observed a correlation between elevated TSSK6 and reduced relapse free survival time exclusively in colon cancer (*P*=0.0092; log rank test; Hazard Ratio (HR), 1.35; 95% confidence interval (*CI*), 1.08-1.69. (Supplemental Figure 1C and Figure 1A). CRC tumors frequently exhibit activating mutations in oncogenes such as *K-RAS* and loss of function mutations in *APC* and *TP53*. HOwevdr, querying TCGA datasets did not reveal strong correlations between TSSK6 expression and mutation of any of these genes (Supplemental Figure 1D). HSP90AB1 interacts with TSSK6 and is frequently upregulated in CRC, however we did not observe a correlation between expression of these two genes in TCGA dataset (Spearman=0.11, p=0.105) (17, 22). To determine the extent of TSSK6 protein accumulation in human colorectal cancer tissue, we developed an immunohistochemical staining protocol in human testis (Figure 1B). Staining was observed in the seminiferous tubule in spermatocytes, spermatids, and Sertoli cells.

**Figure 1.**
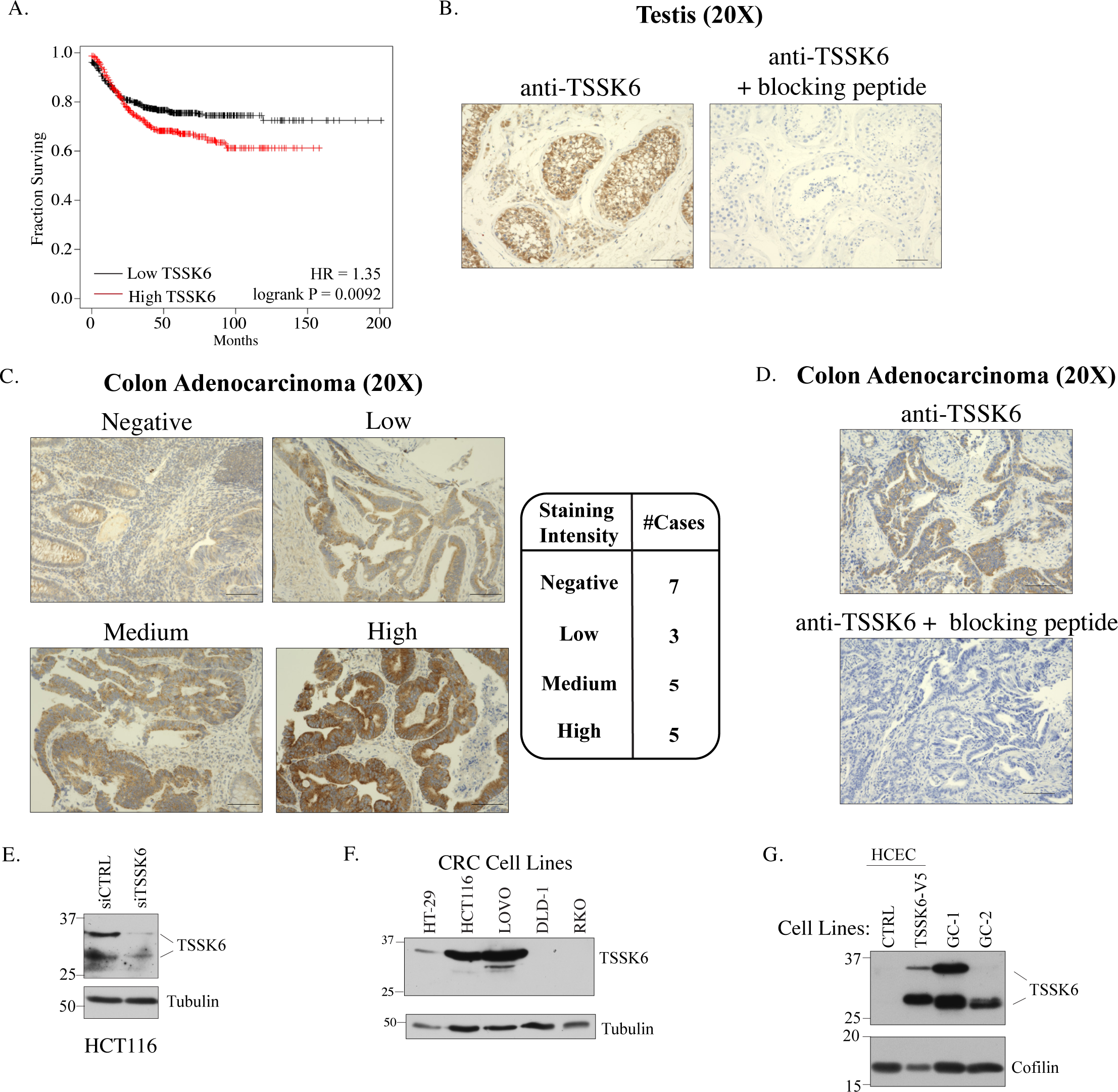
TSSK6 is frequently ectopically expressed in colon cancer. (A) Kaplan Meier curve for relapse free survival based on median expression of TSSK6 in colorectal cancer. (B) Representative images of IHC staining in testis without (left) and with (right) incubation with immunizing peptide. (C). Left: Representative IHC staining of CRC tissues. Scale bars = 100 μm. Right: Associated intensity score for each of the 20 CRC sections stained. (D). Representative staining of CRC section with and without immunizing peptide. Scale bars = 100 μm. (E) HCT116 were transfected with indicated siRNAs for 96 hours, lysed and resolved by SDS-Page (14%) and immunoblotted with indicated antibodies. (F-G) Indicated cell lines were resolved SDS-PAGE (14%) and immunoblotted with indicated antibodies.

Leydig cells of the stroma also exhibited immunoreactivity (Supplemental Figure 2A). The staining was diminished when the sections were co-incubated with the antibody and immunizing peptide (Figure 1B, right panel). Analysis of 20 IHC specimens revealed that ∼65% of CRC tumor cores stained positive for TSSK6 protein with a range of expression levels (Figure 1C). Incubation with a immunizing peptide also fully diminished the TSSK6 signal in CRC specimens (Figure 1D). We next examined endogenous expression of TSSK6 in a human tumor derived CRC cell line, HCT116 (Figure 1E). Here we found that TSSK6 exhibited a slower and faster migrating form on SDS-PAGE, both of which were diminished upon depletion following siRNA targeting TSSK6 (siTSSK6) (Figure 1E). Immunoblots of a larger panel of CRC cell lines, revealed endogenous TSSK6 in HCT116, HT-29 and LOVO CRC cell lines, primarily in the slower migrating form. RKO and DLD-1 cell lines did not express detectable TSSK6 (Figure 1F). We also evaluated expression in a cell line derived from normal human colonic epithelial cells, immortalized with CDK4, hTERT and expressing mutant K-RAS and lacking p53 (referred to as HCECs here) (23). Endogenous TSSK6 was not expressed in these cells, but stable overexpression again resulted in two differentially migrating bands, that were diminished upon siRNA depletion (Figure 1G and Supplemental Figure 2B). Two bands were also detected in immortalized mouse testis cell lines (GC-1, GC-2), which were diminished upon peptide block (Figure 1G and Supplemental Figure 2C). We also detected a band at 100 kDA, but deemed it non-specific as it was not sensitive to TSSK6 siRNA and was present in TSSK6-null cells (Supplemental 2D).

### TSSK6 promotes CRC oncogenicity

We next directly tested whether TSSK6 promotes tumorigenic behaviors. In viability assays, we found that TSSK6 depletion reduced viability of HCT116 cells, but had little impact on HT-29 and LOVO cells (Figure 2A). However, in soft agar assays, depletion of TSSK6 reduced anchorage independent growth in all three TSSK6 expressing lines: HCT116, HT-29 and LOVO (Figure 2B). No effect was observed in TSSK6-negative cell lines (Figure 2B). To control for off-target effects of siRNA, we also verified that a second, independent siRNA pool of two distinct TSSK6 targeting siRNAs recapitulated a defect in anchorage independent growth (Figure 2C). We next measured TSSK6’s contribution to invasion using a transwell assay. We used LOVO cells, which, unlike HCT116, do not exhibit a detectable loss of viability following TSSK6 depletion in 2-D culture (Figure 2A). LOVO cells were depleted of TSSK6 for 48 hours and then measured for invasion over a 48-hour period. In this setting, depletion of TSSK6 was sufficient to dramatically reduce invasive capacity of CRC cells (Figure 2D).

**Figure 2.**
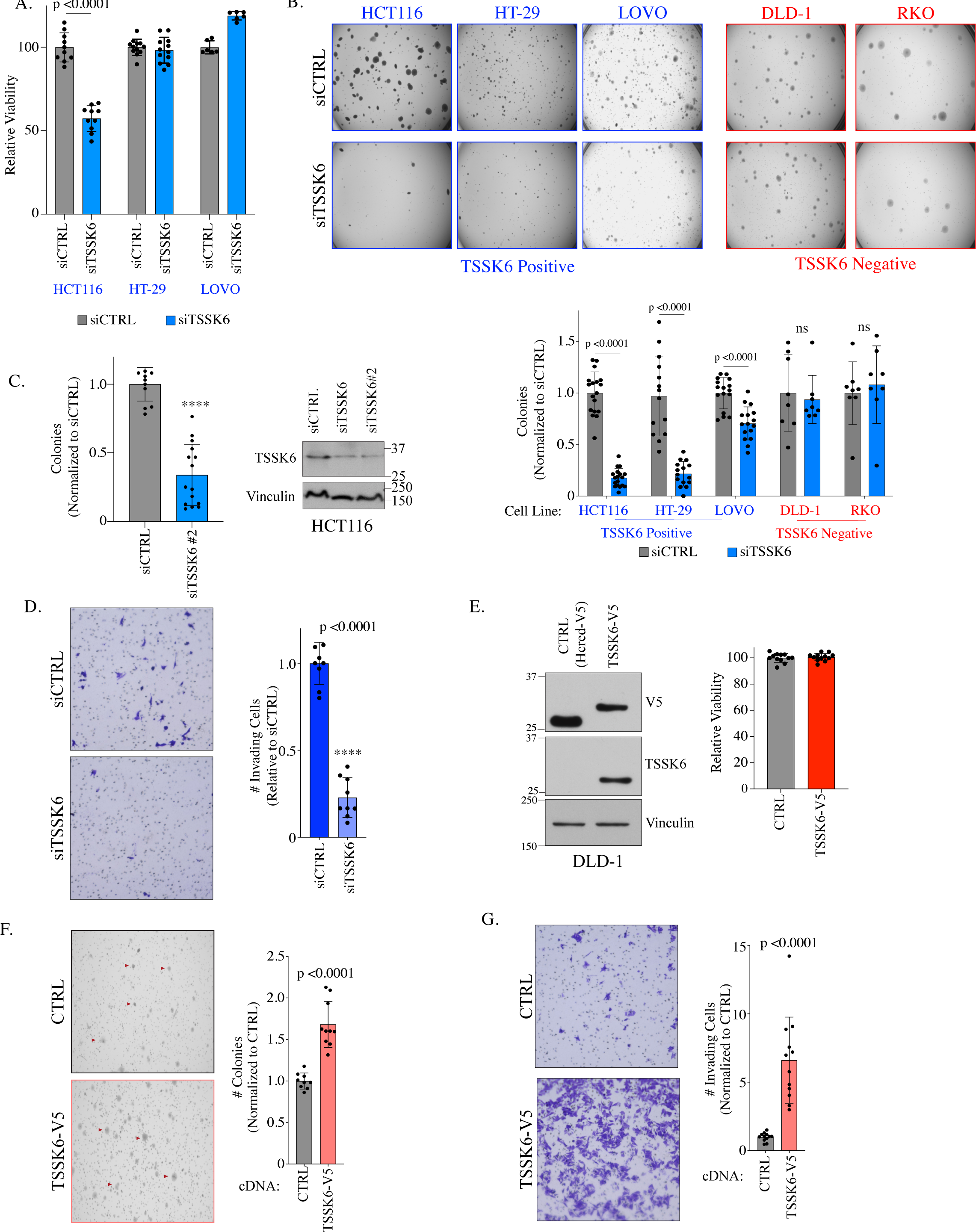
TSSK6 is sufficient for colorectal cancer oncogenic behaviors. (A) Indicated cells were transfected for 96 hours followed by measurement of ATP using Cell Titer Glo. Bars represent mean ± standard deviation (s.d.) (n=6). Statistical analysis by Mann-Whitney t-test. (B) Top: Indicated cell lines were transfected for 48 hours then plated into soft agar. 14-21 days later, samples were stained and quantitated. Below: Quantitation of colonies above. Mean is plotted ± s.d. (ný6). Statistical analysis by Student’s t-test. (C) As in B for second siRNA pool targeting TSSK6 in HCT116 cells. Bars represent mean ± s.d (n=12). Statistical analysis by Mann-Whitney t-test. Right: Parallel lysates from HCT116 cells transfected with two independent siTSSK6 pools were immunoblotted for indicated antibodies.(D) LOVO cells were transfected with indicated siRNA for 48 hours and plated into invasion chambers. 48 hours later, cells were stained and quantitated. Bars represent mean ± s.d. (n=9). Statistical analysis by student’s t-test. (E) Left: Immunoblot of indicated cell lines with indicated antibodies. Samples resolved on 10% SDS-PAGE Right: 4000 cells were seeded for 96 hours followed by Cell Titer Glo assay. Bars represent mean ± s.d. (n=9). (F) Indicated DLD-1 cells were plated into soft agar for 14 days, stained and quantitated. Left: Representative images with arrow heads indicating colonies. Right: Graph represents mean colonies ± s.d. (n=9). Statistical analysis by Mann Whitney t-test. (G) 50,000 DLD1 cells were plated into invasion chambers for 24 hours and subsequently stained and quantitated. Left: Example image. Right: Graphs represent mean ± s.d. (n=9). Statistical analysis by student’s t-test.

To determine if ectopic TSSK6 expression can enhance oncogenic phenotypes, we stably expressed TSSK6 in DLD-1 cells, CRC cells that do not express detectable endogenous TSSK6 (Figure 1F). The DLD-1 TSSK6-V5 cell line did not exhibit a significant increase in cell viability in 2-dimenionsal culture as compared to control (Figure 2E). However, we did observe an increase in soft agar growth and invasion in TSSK6-V5 expressing DLD-1 cells (Figure 2F & 2G). Together this analysis indicates that TSSK6 is sufficient to enhance tumorigenic phenotypes in colorectal cancer cells.

### TSSK6 exhibits bonafide oncogene activity dependent on its kinase activity

We next asked whether ectopic expression of TSSK6 can confer anchorage independent growth to cells that lack this capacity. To this end, we used normal human colonic epithelial cells that are immortalized with hTERT and CDK4, express the oncogene K-RAS (V12) and lack the tumor suppressor, p53 (HCEC1CTRP referred to here as HCEC) (23, 24) (Figure 1G). While these cells exhibit molecular oncogenic changes, they are unable to carry out transformed behaviors including growth in soft agar or growth in xenografts(23). Similar to the DLD-1 cells, overexpression of TSSK6 did not substantially alter viability in 2-dimensional culture (Figure 3A). However, upon stable expression of TSSK6-V5, we observed a robust induction of anchorage independent growth (Figure 3B). Thus, TSSK6 is sufficient to confer soft agar growth in cells that lack this capacity.

**Figure 3:**
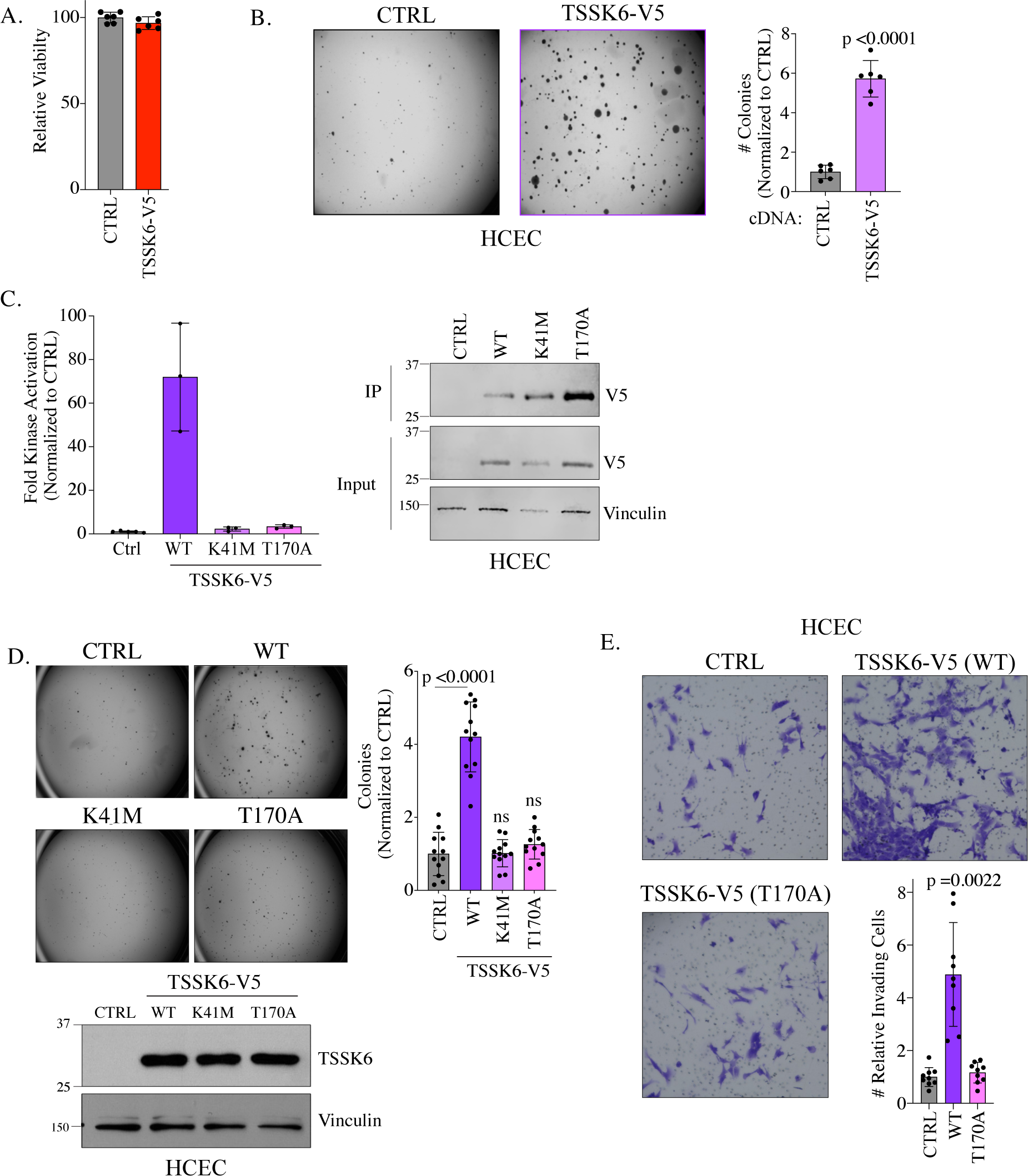
TSSK6 kinase activity is required for transforming activity. (A) Indicated HCEC cell lines were plated for 96 hours and ATP quantitated with Cell Titer Glo. Bars represent mean ± s.d. (n=6). (B) Indicated HCEC cells were grown in soft agar for 21 days then stained and counted. Left: Representative images are shown. Right: Quantitation of colonies. Mean is plotted ± s.d. (n=6). Statistical analysis performed by student’s t-test. (C) Left: Overexpressed TSSK6 was immunoprecipitated with a V5 antibody from HCEC cells and an IP-Kinase assay was used to detect P^32^ incorporation on MBP. Bars represent mean ± s.d. (n=3). Right: Representative blot of parallel lysates resolved on 14% SDS-Page and immunoblotted with indicated antibodies. (D) Indicated HCEC cells were plated into soft agar for 21 days then stained and counted. Left: Representative images. Right: Quantitation of colonies under indicated conditions. Mean is plotted ± s.d. (n=12). Statistical analysis by student’s t-test. Below: Representative immunoblot of indicated cell lines with indicated antibodies. (E) Indicated HCEC cell lines were plated into invasion chambers for 48 hours then stained and counted. Representative images are displayed. Invading cells were quantitated and graph represents mean ± s.d. (n=6). Statistical analysis by Mann Whitney t-test.

TSSK6 is reported to phosphorylate histones during the histone to protamine transition in developing sperm (16). In cells, TSSK6 requires HSP90 for kinase activity. In addition, TSSK6 associates with TSSK6 Activating Co-Chaperone (TSACC, SIP), a tetratricopeptide repeat domain containing protein. TSSK6 recruits TSACC to HSP90 and the formation of this complex leads to maximal TSSK6 activity in 293 cells (21). While HSP90 is widely expressed, TSACC is exclusively expressed in elongated spermatids and not detected in any other normal tissues (21). Thus, we asked whether TSSK6 is an active kinase when ectopically expressed in HCEC cells, outside of its native environment. To measure kinase activity, we adapted a previously published protocol(17). We immunoprecipitated (IPed) stably expressed TSSK6-V5 from HCEC cells and performed an in vitro kinase assay using myelin basic protein (MBP) as a substrate. Wild-type TSSK6 immunoprecipitates exhibited ∼70 fold activity kinase activity over negative control (no TSSK6) (Figure 3C). Mutation of the catalytic lysine (K41M) or the activation loop threonine (T170A), both dramatically reduced kinase activity in this assay (Figure 3C) (21). We subsequently asked whether TSSK6 kinase activity is required for its oncogenic activity. Indeed, HCEC cells expressing wild-type, but not K41M or T170A, TSSK6 exhibited soft agar growth (Figure 3D). In invasion assays, expression of wild-type TSSK6 but not the activation loop mutant, T170A, dramatically increased invasive capacity of HCEC cells (Figure 3E). Together, these findings indicate that TSSK6 kinase activity is required for promoting anchorage independent growth and invasion.

### TSSK6 is associated with and promotes focal adhesion formation

TSSK6 is reported to localize to the nucleus and head and neck region of sperm where it regulates histone phosphorylation and actin polymerization, respectively(17–19). Thus, we asked where TSSK6 localizes when it is inappropriately expressed in cancer cells. In HCT116 cells, we observed endogenous TSSK6 localization throughout the cell, with an enrichment at the nuclear and cellular membrane (Figure 4A). This staining was substantially diminished upon siTSSK6 (Figure 4A). In HCEC cells, ectopically expressed TSSK6-V5 also localized throughout the cell and at the cell membrane where we noted focal enrichment of TSSK6-V5 at the cell membrane (Figure 4B). These foci stained positive for paxillin, a focal adhesion protein (Figure 4B). Moreover, the number of paxillin positive foci were significantly increased in HCEC-TSSK6-V5 cells as compared to HCEC-HcRed cells (Figure 4C). We observed a similar trend for localization and enrichment with tensin, another focal adhesion protein (Figure 4D&E). This finding suggests that TSSK6 promotes focal adhesion formation, potentially through local interactions with proteins that regulate cell-ECM attachments.

**Figure 4.**
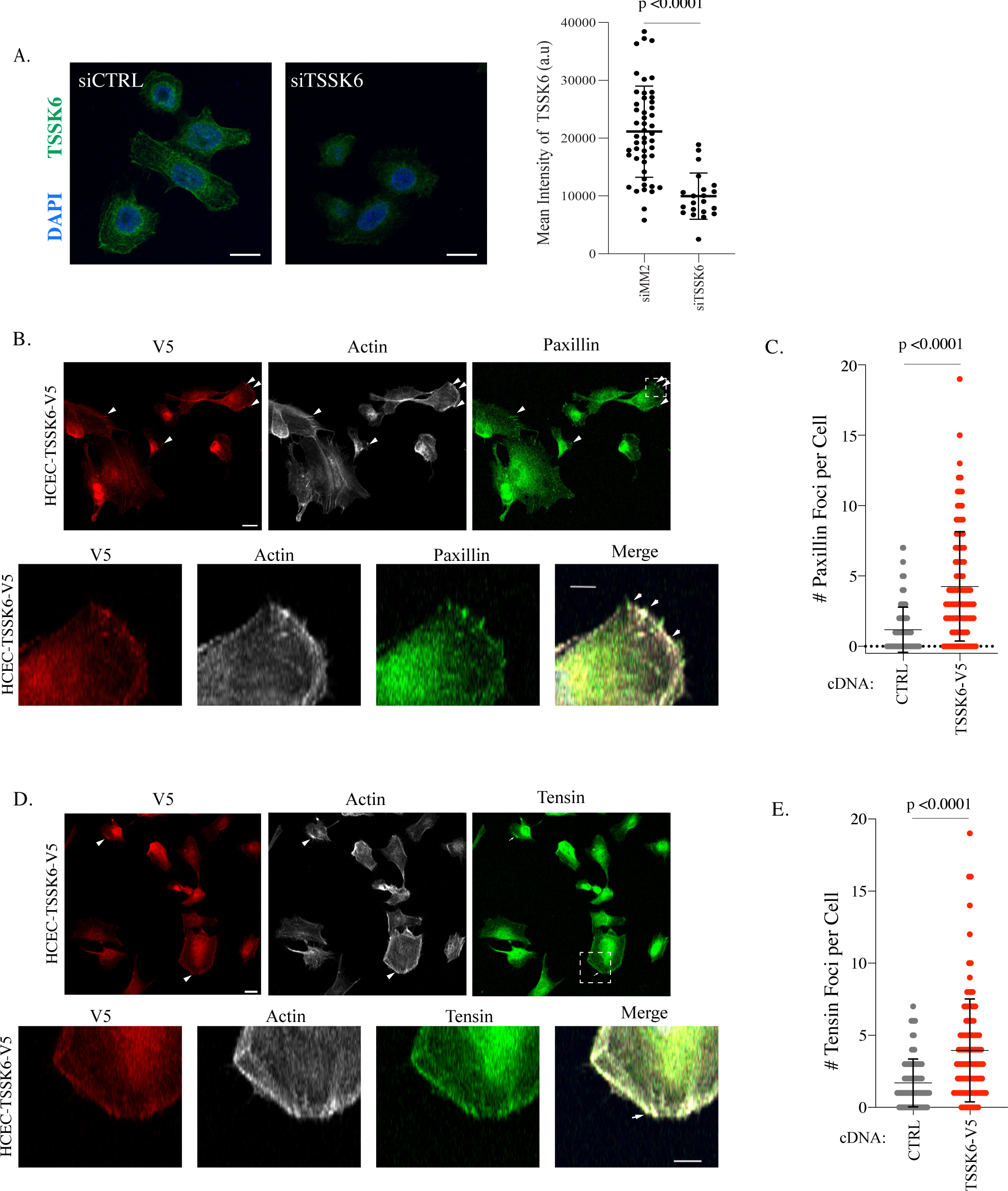
TSSK6 localizes to and enhances focal adhesion formation. (A) Left: Confocal images of HCT116 cells transfected with indicated siRNAs for 72 hours, fixed and immunostained with indicated antibodies. Scale bar is 20 μm. Right: Quantification of TSSK6 signal intensity for individual cells. a.u.= arbitrary units. P-value calculated by student’s t-test (B and C) Left: Confocal images of HCEC cells plated onto glass coverslips for one day prior to staining with indicated antibodies or phalloidin (F-actin). Arrowheads indicate areas of overlap. Scale bar is 20 μm. Hatched box is magnified in lower images with scale bar representing 10 μm. Right: Paxillin and Tensin positive foci were manually quantitated in both the HCEC-Ctrl and HCEC-TSSK6-V5 cells. Mean is indicated with s.d. P-value calculated by Mann-Whitney test.

### TSSK6 promotes tumor growth in vivo

Given the robust in vitro effects observed upon TSSK6 expression, we next examined the impact of TSSK6 on tumor growth in vivo. We established xenograft tumors expressing shCTRL and shTSSK6 in HCT116 cells. We measured tumor growth regularly over 30 days and observed a significant decrease in tumor growth in the mice injected with shTSSK6 cells (Figure 5A). Conversely, we established tumors using DLD-1 cells stably expressing HcRED (Control), TSSK6-wild type or the TSSK6-T170A mutant. Here, we observed enhanced tumor growth in TSSK6-V5 expressing cells as compared to control cells. Notably, TSSK6-V5-170A cells exhibited a nearly identical growth rate to that of control cells, indicating that TSSK6 must be fully active to enhance in vivo growth(Figure 5B, Supplemental Figure 3).

**Figure 5.**
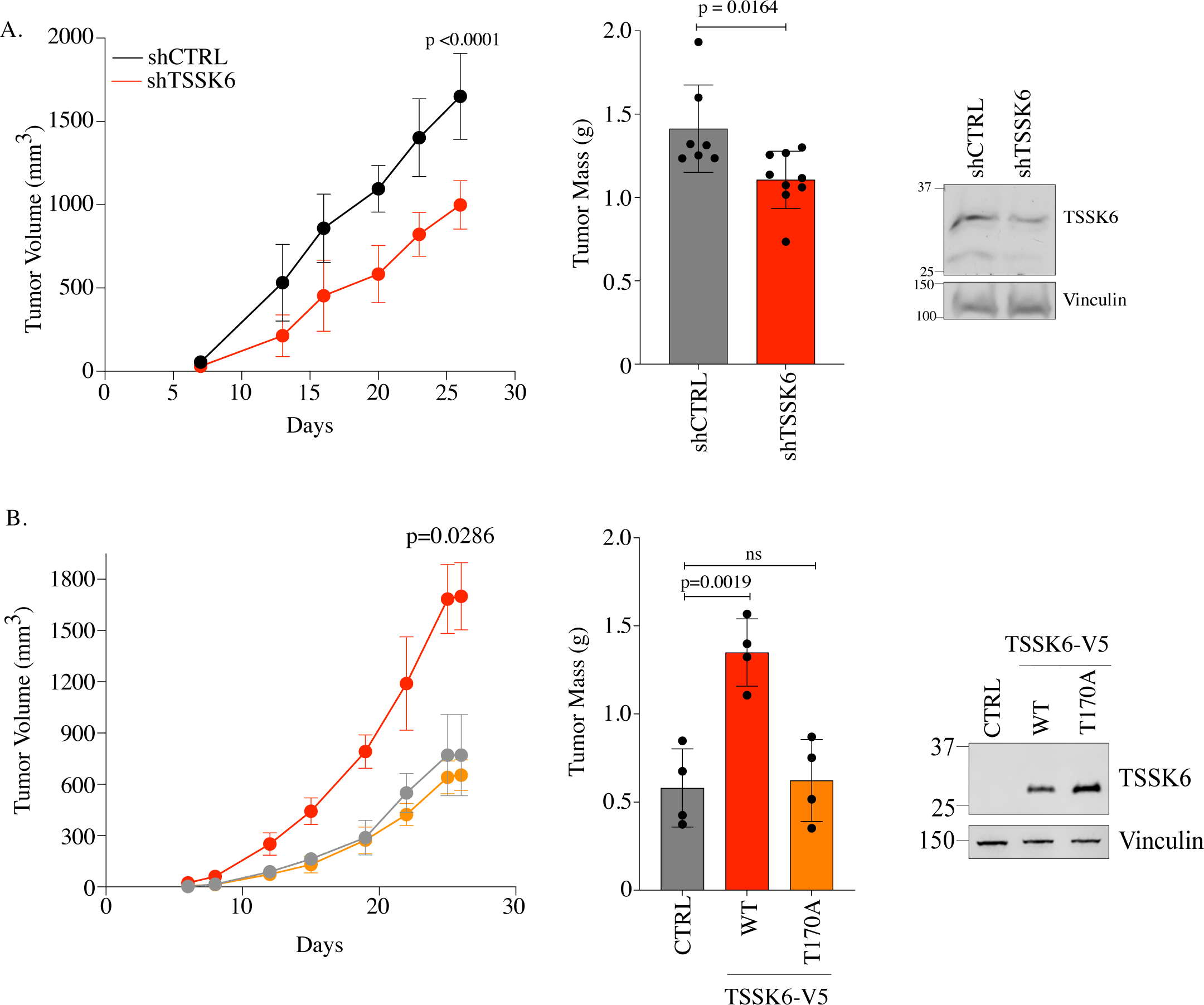
TSSK6 promotes tumor growth in vivo. (A) 1x 10^6^ HCT116 cells stably expressing indicated constructs were xenografted into the flank of NSG mice Left: Tumor volume measurements were taken by caliper on indicated days. Each data point represent the mean (n=9) ± s.d. Middle: Mass of excised tumors. Bars represent the mean (ný7) ± s.d. P-value calculated by Mann Whitney t-test. Upon excision of tumors, two of the largest shCTRL tumors excreted internal contents, thus final masses were not obtainable. Right: Immunoblot of lysates prior to injection resolved on 10% SDS-PAGE and immunoblotted with indicated antibodies. (B) 800,000 DLD-1 cells transfected with indicated constructs were injected into the flank of a NSG mouse. Left: Tumor volume measurements were taken by caliper on indicated days. Each data point represent the mean (n=4) ± s.d. Statistical test performed by Mann Whitney. Middle: Mass of excised tumors. Bars represent the mean. Statistical analysis performed by student’s t-test (n=4). Right: Immunoblot of parallel lysates prior to injection resolved on 10% SDS-PAGE and immunoblotted with indicated antibodies.

## Discussion

Chronic activation of kinases including PIK3CA, EGFR, ALK and RAF are well established to drive tumor progression, survival and metastases. In turn, these kinases present targetable vulnerabilities that when inhibited attenuate tumor growth (25). Our findings here present TSSK6 as a previously unrecognized kinase that promotes oncogenic behaviors. TSSK6 expression is normally restricted to the testes, where it is required for male fertility. Our findings suggest that TSSK6 is frequently aberrantly expressed in colorectal tumors, where it supports tumorigenic behaviors including anchorage independence, invasion and growth in vivo. Based on our findings, we hypothesize that these behaviors are due to TSSK6-induced changes in cell-ECM adhesion properties. These findings suggest the possibility that TSSK6 may confer metastatic capacity to cancer cells, which has been demonstrated in the case of the CTAs, CTAG2 and SPANX(8).

The strong phenotypes induced by TSSK6 expression present a number of implications for future studies. First, it is unknown how TSSK6’s expression and kinase activity are induced when it is abnormally expressed in cancer cells. With respect to expression, CTA promoters are typically highly methylated in somatic cells and DNA methylation can be sufficient to induce CTA expression(26). Whether DNA methylation of TSSK6 is a mechanism for regulating its expression as well as which transcription factors directly activate expression upon demethylation remain unknown. Similarly, mechanisms regulating TSSK6 kinase activity are poorly understood, even in the testes. Studies to-date indicate that HSP70/90 is essential for TSSK6 activation (17, 27). Notably, TSSK6 is inhibited following exposure to the HSP inhibitor 17-allylamino-17-demethoxygeldanamycin (17-AAG) (27). Thus, TSSK6-positive tumors may exhibit enhanced sensitivity to HSP90 inhibitors, which are currently under clinical evaluation in a variety of tumor types (28). TSSK6 activity in sperm is also modulated by the sperm-specific chaperone, TSACC (27). mRNA expression of TSACC is also detectable in tumors and overlaps with TSSK6 in some cases (not shown), suggesting that both proteins may be working in concert in a subset of tumors. Finally, mechanisms that negatively regulate TSSK6 activity in sperm or somatic cells, including phosphatases or degradation pathways have not been explored. If sperm-specific mechanism exists, but are absent in tumors, TSSK6 activity could be relatively unopposed when it is abnormally expressed in somatic cells.

Second, it will be essential to identify TSSK6 substrates and downstream signaling events in cancer cells. A pool of TSSK6 appears to localize to and increase the formation of focal adhesions. Thus, TSSK6 may be regulating cell adhesion through local signaling events at these sites. Studies in sperm and flies, suggest that TSSK6 can regulate cytoskeletal proteins including actin and microtubules, respectively (19, 29). How TSSK6 signaling influences the cytoskeleton and impact cells shape, adhesion and motility are important next questions. The localization of TSSK6 in the cytoplasm, nucleus and at the nuclear periphery also suggest that it shuttles between the cytoplasm and nucleus. TSSK6’s function, if any, in these compartments may also be contributing to its impact on viability and motility of tumor cells. The localization of TSSK6 throughout the cell along with the substantial behavioral changes upon it expression, suggest it could be inducing a multitude of signaling changes that may permit tumor cell adaptation to diverse environments.

Third, our study implicates TSSK6 as a target for anti-cancer therapy. Previous studies have correlated TSSK6 expression with CD8+ T-cell infiltration in multiple tumor types(20). This finding suggests that TSSK6 could generate antigenic peptides in tumors. A number of peptides have been identified with algorithmic tools, but these have not yet been tested for antigenicity (20). Our studies extend the therapeutic potential of TSSK6 and implicate it as a direct interventional target. While TSSK6-specific inhibitors have not been reported, inhibitors have been reported for TSSK2, suggesting this family of kinases is therapeutically tractable (30). If inhibitable with high specificity, TSSK6 could represent an anti-tumor target with an extraordinarily broad therapeutic window.

## Experimental Procedures

### Cell Lines and Culture conditions

GC-1spg and GC-2spd(ts) mouse testis cells were obtained from the American Type Culture Collection (ATCC). HCT116 cells were obtained from Cyrus Vaziri (UNC). RKO, HT-29, DLD-1 and LOVO cell lines were obtained from Jerry Shay (UT Southwestern). Colorectal cancer cells lines were maintained in high glucose DMEM supplemented with 10% FBS (Cat# F0926, Sigma-Aldrich) 1% antibiotic/antimycotic (Cat# 400-101, GeminiBio) and 2.5 mM HEPES pH7.4 at 37°C in a humidified 5% CO_2_ atmosphere. HCEC cells (immortalized with hTERT and CDK4, expressing RASV12 and lacking p53 (HCEC1CTRP)) were developed by Jerry Shay (UT Southwestern) (23, 24). HCECs were maintained in high glucose DMEM supplemented with 5 nM sodium selenite (Cat# S5261, Sigma-Aldrich), 2 ug/mL apo-transferrin (Cat# T1147, Sigma-Aldrich), 20 ng/mL epidermal growth factor (Cat# 236-EG, R&D), 10 ug/mL insulin (Cat# 12585-014, Gibco), 1 ug/mL hydrocortisone (Cat# H0888, Sigma-Alrich), 2% FBS and 1% antibiotic/antimycotic at 37°C in a humidified 5% CO_2_ atmosphere. HEK293T cells were obtained from ATCC and cultured in high glucose DMEM supplemented with 10% FBS, 1% anitbiotic/antimycotic and 2.5 mM HEPES pH7.4 at 37°C in a humidified 5% CO_2_ atmosphere. All cells were periodically evaluated for mycoplasma contamination by a mycoplasma PCR detection kit (Cat# G238, ABM). Authenticity of cell lines was evaluated by detection of ten genetic loci using the GenePrint® 10 System (Promega) and cross referencing to ATCC or internal genetic profiles.

### Antibodies

Antibodies used: TSSK6 (sc-514076, Santa Cruz) (1:100 IHC, 1:500 IB, 1:200 IF), Vinculin (sc-73614, Santa Cruz) (1:2000), Beta-Tubulin (2128, CST) (1:5000), Cofilin (3312, CST) (1:1000), V5-tag (anti-rabbit,13202, CST) (1:5000 IB, 1:500 IF), V5-tag (anti-mouse, 46-0705, Invitrogen)(1.12 ug antibody/500 ug protein IP, 1:2000 IB, 1:100 IF), Paxillin (sc-365379, Santa Cruz) (1:275), Tensin (11990, CST) (1:100).

### Transfections

For siRNA transfections, cells were trypsinized and seeded in Opti-MEM containing Lipofectamine™ RNAiMAX (Thermo Fisher Scientific, Waltham, MA) complexed with siRNAs at a final concentration of 50 nM for all cells except LOVOs, which were transfected with a final concentration of 100 nM. siRNAs for were purchased from Sigma as follows: non-targeting controls (VC30002), targeting TSSK6 pool #1 (referred to as siTSSK6or siTSSK6#1): SASI_Hs01_00190487, (5’CAGUUGCCCUUGUUCGGAA3’) and SASI_Hs01_00190489 (5’AGACAAACUUCUGAGCG 3’). Targeting TSSK6 pool #2 (referred to as siTSSK6#2) WD07275781/82 (5’GCAGCUACUCCAAGGUGAA’3), WD07275783/84(5’GCGUCGUGCUCUACGUCAU 3’).

### Plasmids

cDNAs encoding human *TSSK6* were obtained in pDONR223 (DNASU) and cloned into pLX302 or pLX304 (Addgene Plasmid #25896, #25890) using the Gateway Cloning system (Thermo Fisher Scientific). For control transfections, Hc-Red-V5 in pLX302/pLX304 was used. TSSK6 kinase mutations were made using the QuikChange XL Site-Directed Mutagenesis Kit (200516, Agilent). pLX302-TSSK6-V5 was used as a template. Primers for the K41M mutation were as follows: FWD 5’-CCGTGGCCATCATGGTGGTGGACCG-3’ and REV 5’-CCGTGGCCATCATGGTGGTGGACCG. Primers for the T170A mutation were as follows: FWD 5’-GACCTGAGCACCGCCTACTGCGGCTC-3’ and REV 5’-GAGCCGCAGTAGGCGGTGCTCAGGTC-3’. psPAX2 and pMD2.G lentiviral constructs were purchased from Addgene (Plasmid #12260, #12259). For shRNA experiments, pLKO.1 vectors from TRC expressing TSSK6-targeted shRNAs (TRCN0000199044; sequence: 5’-CGTCGTGCTCTACGTCATGGT-3’, TRCN0000356035 5’-TCGTGCTCTACGTCATGGTCA-3’) were used as shRNA pool. Nontargeting shRNA in pLKO.1 (shSCR) was used as a control (Addgene Plasmid #17920).

### Immunohistochemistry Analysis

Non-malignant testis and colorectal adenocarcinoma tissues were obtained with informed consent from the UTSW Tissue Management Shared Resource (TMSR) in compliance with the UTSW Internal Review Board committee. TSSK6 IHC was optimized and performed by the TMSR according to internal protocols using the Dako Autostainer Link 48 system. Briefly, the slides were baked for 20 minutes at 60°C, then deparaffinized and hydrated before the antigen retrieval step. Heat-induced antigen retrieval was performed at low pH for 20 minutes in a Dako PT Link. The tissue was incubated with a peroxidase block and then an antibody incubated (TSSK6, 1:100, sc-514076, Santa Cruz Biotechnology) for 35 minutes. The staining was visualized using the Keyence BZ-X710 fluorescence microscope. The immunizing peptide (sc-514076 P, Santa Cruz Biotechnology) was incubated at 10:1 mass ratio of peptide:antibody. Staining scores of TSSK6 ranging from 0-3 were set based on the positive scores quantitated by using IHC Profiler of Image J: score 0 (<30), score 1 (30–45), score 2 (45–60), and score 3 (>60) (31). Images were reviewed by a board certified pathologist to validate staining and cellular identity.

### Lentiviral Transduction

Stable cell lines were generated through lentiviral-mediated transduction. HEK293T cells were co-transfected with the target gene vectors and lentiviral packaging plasmids. 48 hours later, virus-conditioned medium was harvested, passed through 0.45 µm pore-size filters, and then used to infect target cells in the presence of Sequabrene™ (S2667, MilliporeSigma) for at least six hours. After infection, medium was replaced, and cells were allowed to recover. DLD-1 cells were infected with pLX304-HcRed-V5 or pLX304-TSSK6-V5 and selected with blasticidin for 3 days or were infected with pLX302-HcRed, pLX302-TSSK6-V5, pLX302-TSSK6-T170A-V5 and selected with puromycin for 3 days. HCEC1CTRP cells were infected with pLX302-HcRed, pLX302-TSSK6-V5, pLX302-TSSK6-T170A, or pLX302-TSSK6-K41M and selected with puromycin for three days. HCT116 cells were infected with pLKO.1 shSCR or shTSSK6 for 120 hours and used immediately for xenograft mouse experiments. Immunoblotting for TSSK6 and/or V5 was used to check for overexpression or depletion.

### Xenograft Assays

All animal experiments were conducted with IACUC approval. 6-10 week old male NOD.cg-PRKDC^SCID^Il2rg^tm1Wjl^/SzJ (NSG) (Rolf Brekken, UT Southwestern) mice were subcutaneously injected in the flank with 1 million cells (HCT116) or 800,000 cells (DLD-1) in 100 µl PBS. Once tumors were visible, tumor volume was measured by caliper in two dimensions, and volumes were estimated using the equation V = length x width^2^/2. Caliper measurements were made twice a week. At the end of the experiment, tumors were harvested and weighed for final tumor mass.

### Immunofluorescence Assays

Cells plated on glass coverslips were washed twice with PBS, fixed with 3.7% formaldehyde for 10 minutes at room temperature. Cells were then washed twice with PBS and incubated in 0.25% Triton-100 for 10 minutes prior to washing three times with 1X PBS. Next coverslips were blocked for at least 30 minutes at room temperature or overnight at 4°C in blocking buffer (1% BSA in 0.001%TBST mixed with PBS). Primary antibodies were diluted with blocking buffer and incubated with samples for 1 hour at room temperature. Next, samples were washed three times in blocking buffer for 5 minutes each, followed by incubation with Alexa Fluor-conjugated secondary antibodies (Thermo Fisher Scientific) in blocking buffer at a dilution of 1:1000 for one hour at room temperature. Phalloidin (1:40) was used to visualize F-actin (Invitrogen, #22287). Lastly, samples were washed three times for five minutes with 0.5% BSA in water and 1 time with water, followed by mounting onto glass slides using ProLong™ Gold Antifade reagent with DAPI. Images were acquired by confocal microscope Zeiss LSM700. Staining was quantitated using Zen 3.8 software (Zeiss). Signal intensity of TSSK6 endogenous staining was quantitated with Zen 3.8 (Zeiss).

### Cell Lysis and Immunoblotting (IB)

Samples were lysed in preheated (100°C for five minutes) 2X Laemmli sample buffer with Beta-mercaptoethanol and boiled for six minutes. Samples were resolved using SDS-PAGE (% for TSSK6 gels noted in legends), and transferred to an Immobilon PVDF membrane (Millipore), blocked in 5% non-fat dry milk followed by incubation in the respective primary antibodies overnight. Immunizing peptide was used at a 5:1 ratio by weight where indicated. Following incubation, membranes were washed three times with Tris Buffered Saline (20 mM Tris, 150 mM NaCl, 0.1% Tween-20) (TBST), and incubated for one hour (overexpressing) to overnight (endogenous) with horseradish peroxidase-coupled secondary antibodies (Jackson Immunoresearch). Subsequently, membranes were washed three times with TBST and then developed using SuperSignal™ West Pico PLUS chemiluminescence substrate (Thermo Fisher scientific, 45-000-875). Immunoblots were scanned using the EPSON perfection v700 photo scanner. For Figure 3C and Figure 5B, after incubation with primary antibodies, membranes were incubated for one hour with fluorophore-coupled secondary antibodies (Licor). Subsequently, membranes were washed three times with TBST and then imaged using Licor Odyssey XF Imaging System.

### Soft agar assays

Cells were suspended into 0.366% bacto-agar and then plated onto solidified 0.5% bacto-agar. Cells were seeded at a density of 500 cells (HCT116 and HCEC1CTRPs), 2000 cells (LOVO, HT29) or 5000 cells (DLD-1 and RKO) per 12-well plate. LOVO, HT29 and HCT116 cells were transfected with siRNA for 48 hours and then collected for plating into soft agar. After 2-3 weeks, colonies were stained for 1 hour with 0.01% crystal violet in 20% methanol. Background stain was removed by washing 3x in water for 30 minutes and 1x overnight. Images were captured with a Leica S9D dissecting microscope, and quantitated by ImageJ software.

### Invasion Assay

Corning BioCoat Growth Factor Reduced Matrigel Invasion Chambers with 8.0 uM PET membranes (Cat# 354483) were used for invasion assays according to the manufacture’s guidelines for use. Briefly, inserts were allowed to come to room temperature for 30 minutes in a sterile hood. Next inserts were allowed to rehydrate in media at 37°C in a humidified 5% CO_2_ atmosphere. After two hours, cell suspensions were prepared at 50,000 cells in 500 uL of 0.1% FBS DMEM per well insert. Media was carefully removed from the invasion inserts and placed into 24-well plates with 10% FBS DMEM. Cell suspensions were immediately added to the wells. Plated inserts were incubated for 24 hours (DLD-1) or 48 hours (HCEC, LOVO). For experiments in which siRNA were used, cells were transfected for 48 h and then collected for plating into transwell inserts.

### Cell Titer Glo Assays (Viability)

Cell-Titer Glo (Promega) (CTG) was performed by manufacture’s protocol but modified to use 15 μl of CTG for 150 μl of cells in media. HCT116, HT29, LOVO cells were reverse transfected with siRNA as indicated and plated at 1000 cells per well and allowed to grow for 96 hours. DLD-1 cells were plated at 4000 cells per well and allowed to grow for 96 hours. HCECs were plated at 1000 cells per well and allowed to grow for 96 h. Luminescence was read using a CLARIOstar Plus plate reader (BMG Labtech).

### Immunoprecipitation(IP) -Kinase assays

Cells were lysed for 30 minutes on ice in non-denaturing lysis buffer (50 mM HEPES pH 7.4, 50 mM NaCl, 0.1 % Triton X-100, 100 mM NaF, 30 mM sodium pyrophosphate, 50 mM ϕ3-glycerophosphate 1 mM EGTA, 10% glycerol, 1 mM Na_3_VO_4_, 0.4 ug/mL pepstatin, 0.4 ug/mL leupeptin, 4 ug/mL TAME, 4 ug/mL tos-lys-chloromethylketone, 4 ug/mL Na-benzoyl-L-arginine methyl ester carbonate, 4 ug/mL soybean trypsin inhibitor). Lysates were clarified at 12,000 x g for 10 minutes and pre-cleared with Protein A/G agarose beads for one hour at 4°C. Five percent of each lysate was set aside as input material and the remainder of lysate was incubated with 1.12 ug of V5 antibody and for one hour at 4°C. 35 ul of protein A/G agarose beads were then added and incubated for 1 hour at 4°C. Beads were washed three times with 1 mL of 1M NaCl, 20 mM Tris pH 7.4 and one time with minimal kinase reaction buffer (10 mM HEPES pH 8.0, 10 mM MgCl2). Kinase assays were carried out at 30 °C for 30 minutes in 10 mM HEPES pH 8.0, 10 mM MgCl2, 1 mM benzamidine, 1 mM DTT, 50 μm ATP (1 cpm/fmol ATP-P^32^) with 2 uL 5mg/mL Myelin Basic Protein (MBP) as a substrate. Reactions were stopped by adding 10 μl of GenScript 4X LDS sample buffer, followed by boiling for 3 minutes. Samples were run on GenScript 4-20% Sure Page gels. Gels were stained with coomassie blue and bands corresponding to MBP were excised and P^32^ incorporation measured using a liquid scintillation counting. These values were then normalized to control immunoprecipitates to obtain a fold activation. This protocol is adapted from previously described assays measuring TSSK6 and ERK2 phospho-transfer activity. (17, 32)

### Normal and tumor expression analysis

The Gtex Portal was used to analyze TSSK6 mRNA expression across normal tissues on 12/2023. The Genotype-Tissue Expression (GTEx) Project was supported by the Common Fund of the Office of the Director of the National Institutes of Health, and by NCI, NHGRI, NHLBI, NIDA, NIMH, and NINDS. cBIO Portal was used to analyze expression in tumors in the TCGA PanCancer Atlas Study (10,528 cases) using a z-score cutoff of 2 (21).

### Survival Analysis

Survival analysis was performed using the KM Plotter program (33). Affymetrix 224409_s_t was used for survival analysis for TSSK6. Populations were split based on median expression and relapse-free and overall survival was measured. Analysis was completed in December of 2023.

### TCGA Data Analysis

Using the Firehose Legacy TCGA Data set in cBioportal, we extracted colorectal cancer cases with micoarray expression analysis and sequencing of KRAS, TP53 and APC available (n=205). TSSK6 expression was ranked by z-score. KRAS, TP53 and APC mutation status was extracted for each case in the top and bottom quartiles of TSSK6 expression (n=51 cases each). A chi-squared test (Graphpad Software) was used to compare incidence of mutation of each individual gene in the top and bottom quartiles TSSK6 populations. cBio Portal was used to examine expression correlation between TSSK6 and HSP90AB1 using the Firehose legacy expression data from microarray analysis (21).

### Statistical Analysis

Graphpad Prism (Graphpad Software) was used to perform statistical analyses. Normality of data was determined by the Shapiro-Wilk normality test. Statistical differences in all cases were determined by Mann-Whitney (not normal distribution) or t-test (normal distribution) as specified in the figure legends. : ns= not significant, P > 0.05; *P < 0.05; **P < 0.01; ***P < 0.001; ****P < 0.0001. Outliers were detected by ROUT analysis.

## Author contributions

A.W.W., M.D. M.H.C. conceptualization; A.W.W., M.D., Z.G., methodology; A.W.W.,Z.G.,K.M., S.S., M.D., investigation; A.W.W. and M.D. writing—original draft, review & editing; A.W.W., supervision; A.W.W. funding acquisition.

## Funding

Funding was provided by the National Institutes of Health to A.W.W. (R01CA251172 and R01CA196905, R03CA250021). M.D. was supported by the Cancer Prevention Research Institute of Texas (CPRIT) Training Grant (RP210041) and a T32 NCI Training Grant (CA124334-15). This work was also supported by The Welch Foundation to A.W.W (I-2087-20210327). Support for use of core services was made possible through NCI Cancer Center Support Grant (5P30CA142543) to the Harold C. Simmons Cancer Center. The content is solely the responsibility of the authors and does not necessarily represent the official views of the National Institutes of Health.

## Conflict of Interest

The authors declare that they have no conflict of interest

## Acknowledgements

The authors would like to thank Jerry Shay for the HCEC cells, and Michael Reese and Melanie Cobb for helpful discussions. The authors also thanks Diego Castrillon for reviewing pathology samples and Cheryl Lewis for IHC staining and tissue procurement.

## Data Availability

All data is contained within the manuscript

## Notes

### Competing Interest Statement

The authors have declared no competing interest.

### Summary of Updates

Added supplemental figures and updated the text.

