## Supplemental Figures for "Testis Specific Serine Kinase 6 (TSSK6) is abnormally expressed in colorectal cancer and promotes oncogenic behaviors"

Supplemental Figure Legend:

Figure S1: A. Graph of TSSK6 mRNA expression across human tissues derived from the GTex database. B. Oncoprint of TSSK6-expressing tumors (pink) with tumor type indicated based on TCGA PanCancer Atlas Study. C. Patient outcome data for indicated tumors based on TSSK6 mRNA median expression. D. TCGA CRC cases exhibiting the top and bottom quartile of TSSK6 expression and associated mutation of indicated genes. P-value calculated by chi-squared analysis (Prism Graphpad) Software.

Figure S2: A. Zoomed cross section of human testes tissue stained with anti-TSSK6. Red labelling indicates TSSK6-positive cells and black labelling indicates TSSK6-negative cells. B. Indicated HCEC lysates resolved by SDS-PAGE (10%) and immunoblotted with anti-TSSK6. C. Mouse spermatogonial cell lysates were resolved by SDS-PAGE (14%) and membranes incubated +/- immunizing peptide. D. Full blot image of indicated lysates resolved on 14% SDS-PAGE and probed with a anti-TSSK6 antibody. Arrow indicates non-specific band at ~100kDa.

Figure S3: DLD-1 xenograft tumors +/- V5-TSSK6 (Figure 5B) were harvested, fixed and stained using the TSSK6 IHC protocol. Scale bar represents 100 um.

Delgado et al
Supplemental Figure 1

A.

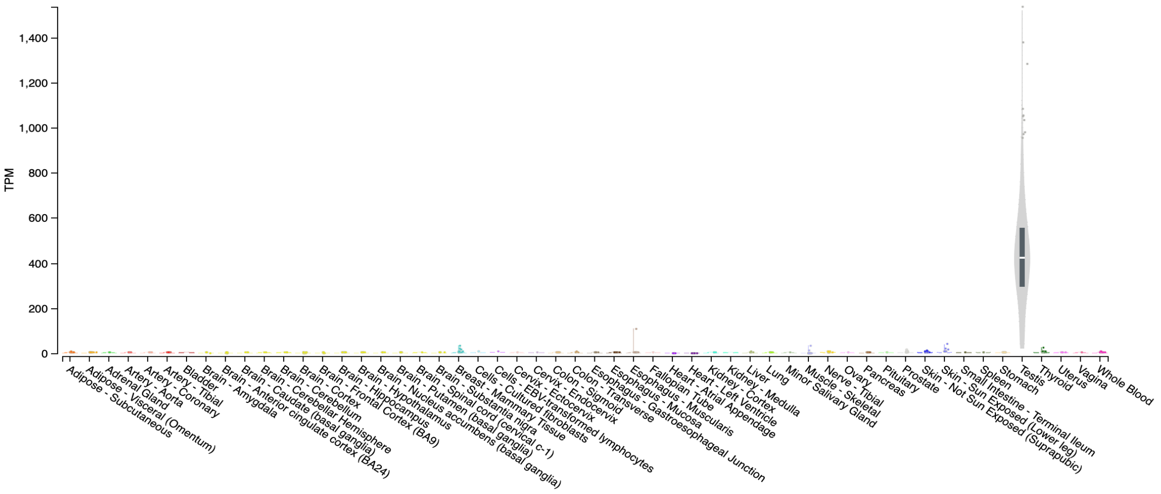


B.

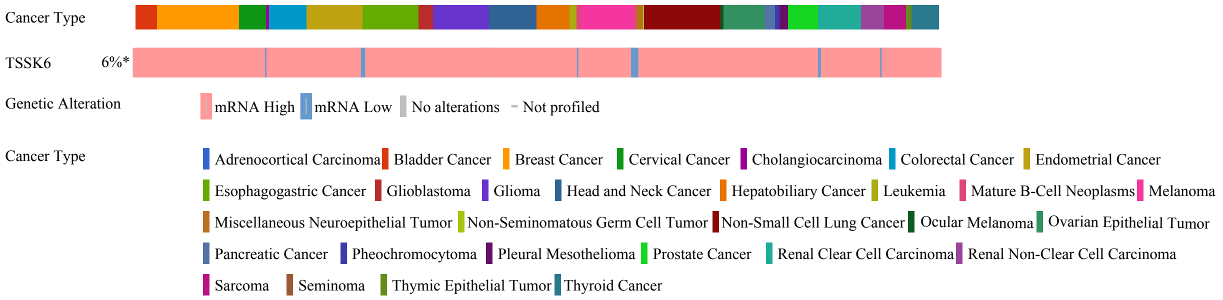


C.

| Tumor Type | Survival | # Patients | Hazard Ratio | p-Value | Survival | # Patients | Hazard Ratio | p-Value |
|---|---|---|---|---|---|---|---|---|
| Breast | RFS | 2032 | 0.99 | 0.92 | OS | 943 | 1.13 | 0.38 |
| Ovarian | RFS | 614 | 0.95 | 0.5609 | OS | 655 | 0.99 | 0.56 |
| Lung | FP | 874 | 1.22 | 0.074 | OS | 1411 | 0.79 | 0.003 |
| Gastric | FP | 522 | 1.06 | 0.63 | OS | 631 | 1.22 | 0.074 |
| Pancreatic | DFS | 278 | 1.1 | 0.435 | OS | 1189 | 0.9 | 0.128 |
| **Colorectal** | **RFS** | **1130** | **1.35** | **0.0092** | OS | 809 | 1.11 | 0.39 |

RFS: Relapse Free Survival
DFS: Disease Free Survival
FP: Time to First Progression
OS: Overall Survival

D.

| KRAS | | | |
|---|---|---|---|
| TSSK6 population: | Mutant # cases | Wild-type # cases | chi-square p-value |
| Top quartile | 18 | 33 | 0.0727 |
| Bottom quartile | 27 | 24 | |

| APC | | | |
|---|---|---|---|
| TSSK6 population: | Mutant # cases | Wild-type # cases | chi-square p-value |
| Top quartile | 33 | 18 | 0.8368 |
| Bottom quartile | 32 | 19 | |

| p53 | | | |
|---|---|---|---|
| TSSK6 population: | Mutant # cases | Wild-type # cases | chi-square p-value |
| Top quartile | 29 | 22 | 0.2347 |
| Bottom quartile | 23 | 28 | |

Delgado et al
Supplemental Figure 2

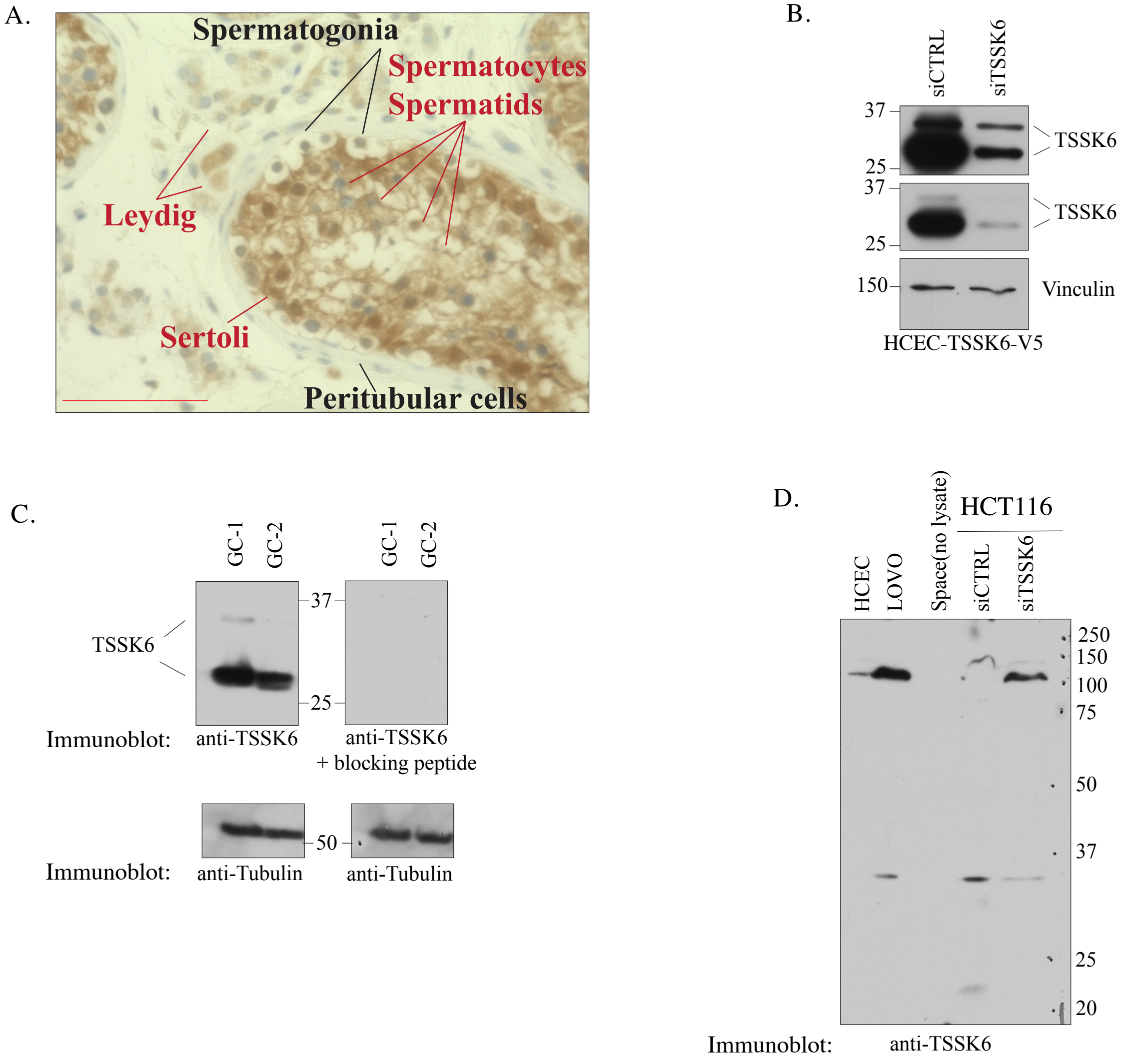

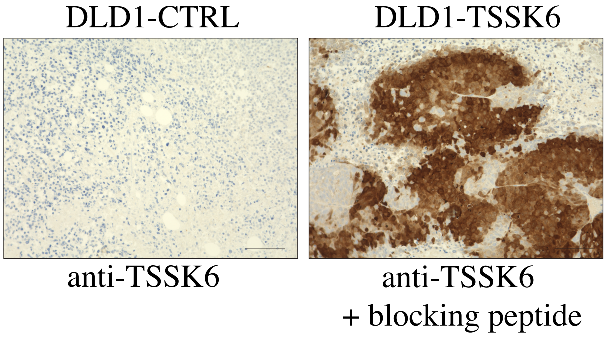
DLD1-CTRL
DLD1-TSSK6
anti-TSSK6
anti-TSSK6
+ blocking peptide
